# Defects in Meiotic Recombination Delay Progression Through Pachytene in Mouse Spermatocytes

**DOI:** 10.1101/102251

**Authors:** James H. Crichton, David Read, Ian R. Adams

## Abstract

During meiosis, recombination, synapsis, chromosome segregation and gene expression are coordinately regulated to ensure successful execution of this specialised cell division. In many model organisms, checkpoint controls can delay meiotic progression to allow defects or errors in these processes to be repaired or corrected. Mouse spermatocytes possess quality control checkpoints that eliminate cells with persistent irreparable defects in chromosome synapsis or recombination, and here we show that a spermatocyte checkpoint regulates progression through pachytene to accommodate delays in meiotic recombination. We have previously show that the appearance of early recombination foci is delayed in *Tex19.1*^*-/-*^ spermatocytes during leptotene/zygotene, but some *Tex19.1*^*-/-*^ spermatocytes still successfully synapse their chromosomes. Therefore, we have used autosomally synapsed *Tex19.1*^*-/-*^ mouse spermatocytes to assess the consequences of delayed recombination on progression through pachytene. We show that these pachytene spermatocytes are enriched for early recombination foci. This skew is not accompanied by cell death and likely reflects delays in the generation and/or maturation of recombination foci. Moreover, patterns of axis elongation, chromatin modifications, and histone H1t expression are also all skewed towards earlier substages of pachytene suggesting these events are co-ordinately regulated. Importantly, the delay in histone H1t expression in response to loss of *Tex19.1* does not occur in a *Spo11* mutant background, suggesting that histone H1t expression is being delayed by a recombination-dependent checkpoint. These data indicate that a recombination-dependent checkpoint operates in mouse spermatocytes that can alter progression through pachytene to accommodate spermatocytes with some types of recombination defect.

## Author Summary

Meiosis is a specialised cell division that reduces the number of chromosomes in each cell during sperm and egg development. A number of processes need to occur in the correct order for meiosis to be successful, and in many species checkpoint mechanisms monitor these processes for defects and can delay meiotic progression until they are repaired. If cells have more persistent or irreparable defects then the same checkpoints can eventually trigger apoptosis and eliminate these cells from the population. Mouse spermatocytes possess pachytene checkpoints that eliminate spermatocytes with severe defects in recombination and chromosome synapsis. Here we show that mouse spermatocytes also possess a meiotic checkpoint that can alter progression through pachytene in response to delayed meiotic recombination. The ability of mouse spermatocytes to delay aspects of progression through pachytene in response to these defects demonstrates that there is some plasticity in meiotic progression that could allow some recombination defects to be repaired. The existence of a meiotic checkpoint that can delay progression of mammalian spermatocytes through pachytene may be particularly important in human populations where mutations, polymorphisms and genetic variation present in natural populations could often result in altered timing or suboptimal execution of some meiotic processes.

## Introduction

Meiosis is a central feature of the life cycle of sexually reproducing organisms that requires coordinated regulation of multiple distinct processes including recombination, chromosome synapsis, chromosome segregation and changes in gene expression, to generate haploid germ cells [1,–3]. The correct temporal execution of some of these processes relies on substrate-product relationships between them, for example meiotic recombination generates aligned homologous chromosomes that are substrates for chromosome synapsis. In addition feedback and feedforward loops help to ensure robust execution of some meiotic processes [4]. One of the feedback systems operating in mouse spermatocytes restricts DNA double strand break (DSB) formation to the asynapsed regions of chromosomes [5]. Evidence for this feedback system can be seen in spermatocytes that have reduced activity of SPO11, a subunit of the endonuclease that generates meiotic DSBs [5]. These hypomorphic *Spo11* spermatocytes generate fewer DSBs during leptotene causing reduced homologous chromosome synapsis, but a feedback mechanism allows the resulting asynapsed chromosomal regions to accumulate high densities of DSBs during late zygotene, presumably in order to stimulate the homology search in these regions and promote their synapsis [5].

In addition to feedback controls, checkpoints are also an important component of meiotic regulation. Checkpoints monitor and co-ordinate meiotic events, and can provide a quality control mechanism to eliminate cells that do not execute some aspects of meiosis correctly [4]. Persistent defects in chromosome synapsis in mouse spermatocytes can activate a checkpoint that triggers cell death during pachytene [6]. This synapsis checkpoint is caused by asynapsed chromosomes sequestering the transcriptional silencing machinery away from the heterologous sex chromosomes, causing defective meiotic sex chromosome inactivation (MSCI) and inappropriate expression of sex chromosome-encoded gene products [7]. Moreover, oocytes and spermatocytes also possess meiotic checkpoints that trigger prophase cell death in response to recombination defects, although there are differences in the signalling pathways involved between the sexes [8–10]. The tight coupling between recombination and synapsis means that many meiotic recombination mutants activate the synapsis checkpoint in spermatocytes [6], making it difficult to study the effects of perturbed recombination on meiotic progression in the absence of confounding effects of asynapsis. However, a recombination checkpoint has been proposed to operate in spermatocytes carrying moderate severity mutations in *Trip13 [10]*. *Trip13* encodes an AAA+ ATPase implicated in regulating HORMAD proteins [11], and moderate severity *Trip13* mutants successfully synapse their chromosomes but have defects in the maturation of recombination foci [12,13]. These spermatocytes accumulate markers of unrepaired DNA damage and die through activation of a DNA damage checkpoint [10,12,13].

In addition to chromosome synapsis and DNA repair checkpoints operating in prophase, a spindle assembly checkpoint can delay the onset of anaphase I in response to defects in chromosome alignment in mouse meiosis [14]. This checkpoint appears to be more sensitive in spermatocytes than oocytes [15,16]. Furthermore, even when this checkpoint is activated, mouse oocytes with misaligned chromosomes can eventually proceed into anaphase I after a delay, whereas similar defects induce a strict metaphase I arrest and cell death in spermatocytes [17,18]. Thus, checkpoint activation can delay cell cycle progression to allow the cells to repair defects before progressing to the next stage of the cell cycle, and cells with persistent irreparable defects eventually continue through the checkpoint or initiate cell death depending on the context [4,19]. Although the pachytene asynapsis and DNA damage checkpoints have primarily been demonstrated to cause apoptosis and meiocyte death in mice, it is not clear whether these checkpoints are also able to coordinate meiotic progression to accommodate suboptimal cells that have encountered reparable lesions or delays.

*Tex19.1* was originally isolated as a testis-expressed gene, and encodes a mammal-specific protein that interacts with the E3 ubiquitin ligase UBR2 [20,21]. *Tex19.1* plays a role in repressing retrotransposon expression in germ cells and hypomethylated somatic tissues [22,23], but potentially has additional targets that cause defects in meiotic recombination [24]. We have recently shown that *Tex19.1*^*-/-*^ spermatocytes have reduced numbers of early recombination foci during leptotene and that, similar to hypomorphic *Spo11* spermatocytes [5], additional early recombination foci are generated during zygotene in these mutants [24]. This early recombination defect likely causes the defects in homologous chromosome synapsis seen around two-thirds of pachytene *Tex19.1*^*-/-*^ spermatocytes [24]. However, the remaining one-third synapse their autosomes completely. These autosomally synapsed *Tex19.1*^*-/-*^ spermatocytes therefore provide a system to study how delays in recombination might influence progression through pachytene in the absence of the confounding effects of asynapsis.

Here we show that progression through pachytene is perturbed in autosomally synapsed *Tex19.1*^*-/-*^spermatocytes. We show that the progression, timing or duration of recombination, chromosome axis elongation, chromatin modifications, and histone H1t expression are co-ordinately altered in *Tex19.1*^*-/-*^ spermatocytes, and that the altered progression of *Tex19.1*^*-/-*^ spermatocytes through pachytene depends on *Spo11*. These findings indicate that meiotic recombination is monitored by a checkpoint in spermatocytes that can delay aspects of progression through pachytene to accommodate cells with altered recombination kinetics.

## Results

### Early Recombination Markers Are Enriched In Autosomally Synapsed *Tex19.1*^*-/-*^ Pachytene Spermatocytes

We have previously shown that *Tex19.1*^*-/-*^ spermatocytes have defects in the initiation of *Spo11*-dependent early recombination foci [24]. The number of DMC1-containing recombination foci in *Tex19.1*^*-/-*^ spermatocytes is reduced to around 30% of those present in littermate controls during leptotene, but increases to reach 87% of control levels during zygotene [24]. Thus, a significant number of recombination foci are being generated during zygotene rather than leptotene in *Tex19.1*^*-/-*^ spermatocytes. These early recombination defects are accompanied by synapsis defects in some *Tex19.1*^*-/-*^ pachytene spermatocytes, but the remaining fully synapsed pachytene spermatocytes must have possessed sufficient recombination foci to promote a successful homology search, and do not have any confounding autosomal asynapsis that would interfere with analysing the effects of delayed recombination on meiotic progression. We have therefore used these autosomally synapsed pachytene *Tex19.1*^*-/-*^ nuclei to investigate whether delayed recombination might subsequently affect progression through the pachytene stage of meiosis.

To investigate the kinetics of maturation of recombination foci in autosomally synapsed pachytene *Tex19.1*^*-/-*^ spermatocytes, we immunostained chromosome spreads from testes from adult *Tex19.1*^*-/-*^ mice, and from *Tex19.1*^*+/±*^ littermate controls, with antibodies to DMC1, RPA and RAD51 (Figure 1A-1C). DMC1 and RAD51 mark early recombination foci and dissociate as recombination foci mature during pachytene [25]. RPA continues to mark recombination foci after DMC1 and RAD51 dissociate, but is also eventually lost during pachytene as recombination foci mature [25]. Pachytene nuclei with complete autosomal synapsis were identified by immunostaining for synaptonemal complex components, and autosomally asynapsed pachytene nuclei, which are abundant in *Tex19.1*^*-/-*^ testes, were excluded from this analysis. Control pachytene *Tex19.1*^*+/±*^ spermatocytes had 86±5 RPA foci, 13±1 DMC1 foci and 13±2 RAD51 foci on their autosomes (Figure 1D), comparable to other studies [13,26]. Interestingly, autosomal RPA and DMC1 foci in autosomally synapsed *Tex19.1*^*-/-*^ pachytene spermatocytes were significantly increased by 28% and 95% respectively (Figure 1D). Violin plots of these data (Supporting Figure S1) suggest bimodal distributions for both RPA and DMC1, with one population of pachytene nuclei containing relatively high numbers of RPA or DMC1 foci, and a second population containing few or no RPA or DMC1 foci. The increase in RPA foci number in autosomally synapsed *Tex19.1*^*-/-*^ pachytene spermatocytes appears to primarily reflect a shift towards the abundant foci population, whereas the increase in DMC1 foci number reflects both a shift towards the abundant foci population and an increase in the number of DMC1 foci in that population (Supporting Figure S1). RAD51 foci were not significantly increased in *Tex19.1*^*-/-*^ spermatocytes (Figure 1D), although it is not clear if this reflects a genuine difference between RAD51 and its meiotic homolog DMC1 rather than technical differences in antibody sensitivity. Regardless, the increased frequency of DMC1 and RPA foci in pachytene *Tex19.1*^*-/-*^ spermatocytes indicates that, despite successful autosomal synapsis, these cells possess an abnormally high frequency of recombination foci participating in early stages of meiotic recombination.

**Figure 1.**
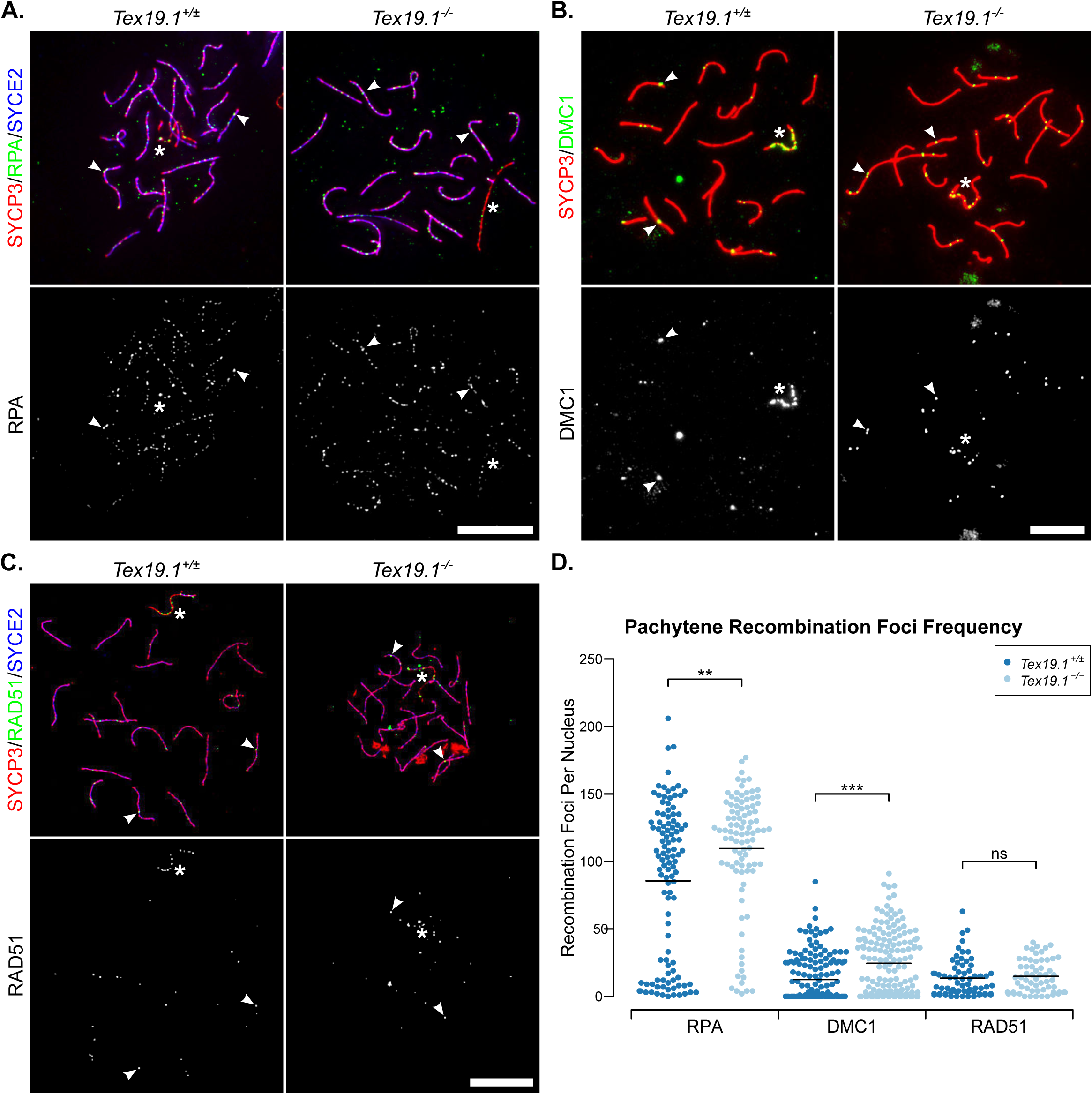
**Autosomally Synapsed Pachytene *Tex19.1***^***-/-***^**Spermatocytes Have Increased Numbers of Early Meiotic Recombination Foci.** (A-C) Immunostaining for RPA (A), DMC1 (B) and RAD51 (C) recombination proteins in *Tex19.1*^*+/±*^and *Tex19.1*^*-/-*^autosomally synapsed pachytene spermatocyte chromosome spreads. The recombination protein are shown in green, SYCP3 (A, B, C, red) and SYCE2 (A, C, blue) mark the lateral and central elements of the synaptonemal complex respectively. Single channel greyscale images for the recombination foci are also shown, foci co-localising with axes were scored as recombination foci. Asterisks indicate sex body, arrowheads indicate example recombination foci. Scale bars 10 μm. (D) Scatterplots showing the number of axial recombination foci in autosomally synapsed pachytene nuclei. Means are indicated by horizontal lines. Mean foci frequencies are 86±5, 110±5, 13±1, 24±2, 13±2, 15±2 from left to right across the plot. Number of nuclei analysed were 109, 94, 165, 163, 66, 60 from a total of at least 3 experimental or 3 control animals for each recombination protein. Foci counts were compared between genotypes using a Mann-Whitney U test, asterisks denote significance (** p < 0.01, *** p < 0.001), ns indicates no significant difference (p > 0.05).

To verify our analysis of early recombination intermediates in pachytene *Tex19.1*^*-/-*^ spermatocytes we examined the behaviour of γH2AX (Figure 2A), a marker for unrepaired DNA damage [27,28]. In pachytene, γH2AX is typically detected as a strong cloud of staining over the asynapsed sex chromosomes, and as condensed axial foci (S-foci) on autosomes that co-localise with recombination foci [28]. The number of autosomal γH2AX S-foci decreases during pachytene and are undetectable in diplotene [28]. As distinguishing individual S-foci becomes somewhat subjective when there are large numbers of adjacent γH2AX S-foci on the same axis, we grouped nuclei as having abundant (> 40), intermediate (1-40) or no γH2AX S-foci. In control mice, 22% of autosomally synapsed pachytene spermatocytes had abundant, 62% had intermediate, and 16% had no γH2AX S-foci (Figure 2B). Autosomally synapsed pachytene spermatocytes with abundant γH2AX S-foci were ~2-fold more frequent in *Tex19.1*^*-/-*^ mice, and correspondingly fewer autosomally synapsed *Tex19.1*^*-/-*^ pachytene nuclei had intermediate or no γH2AX S-foci (Figure 2B). Loss of *Tex19.1* does not have any detectable effect on the frequency of *Spo11*-independent γH2AX foci in spermatocytes [24], suggesting that the increase in the number of pachytene spermatocytes with abundant γH2AX S-foci represents a difference in behaviour of *Spo11*-dependent meiotic DSBs in these mutants. Therefore, in addition to containing elevated levels of early recombination foci, autosomally synapsed *Tex19.1*^*-/-*^ pachytene spermatocytes more frequently contain abundant γH2AX S-foci suggesting there is more unrepaired DNA damage in these nuclei.

**Figure 2.**
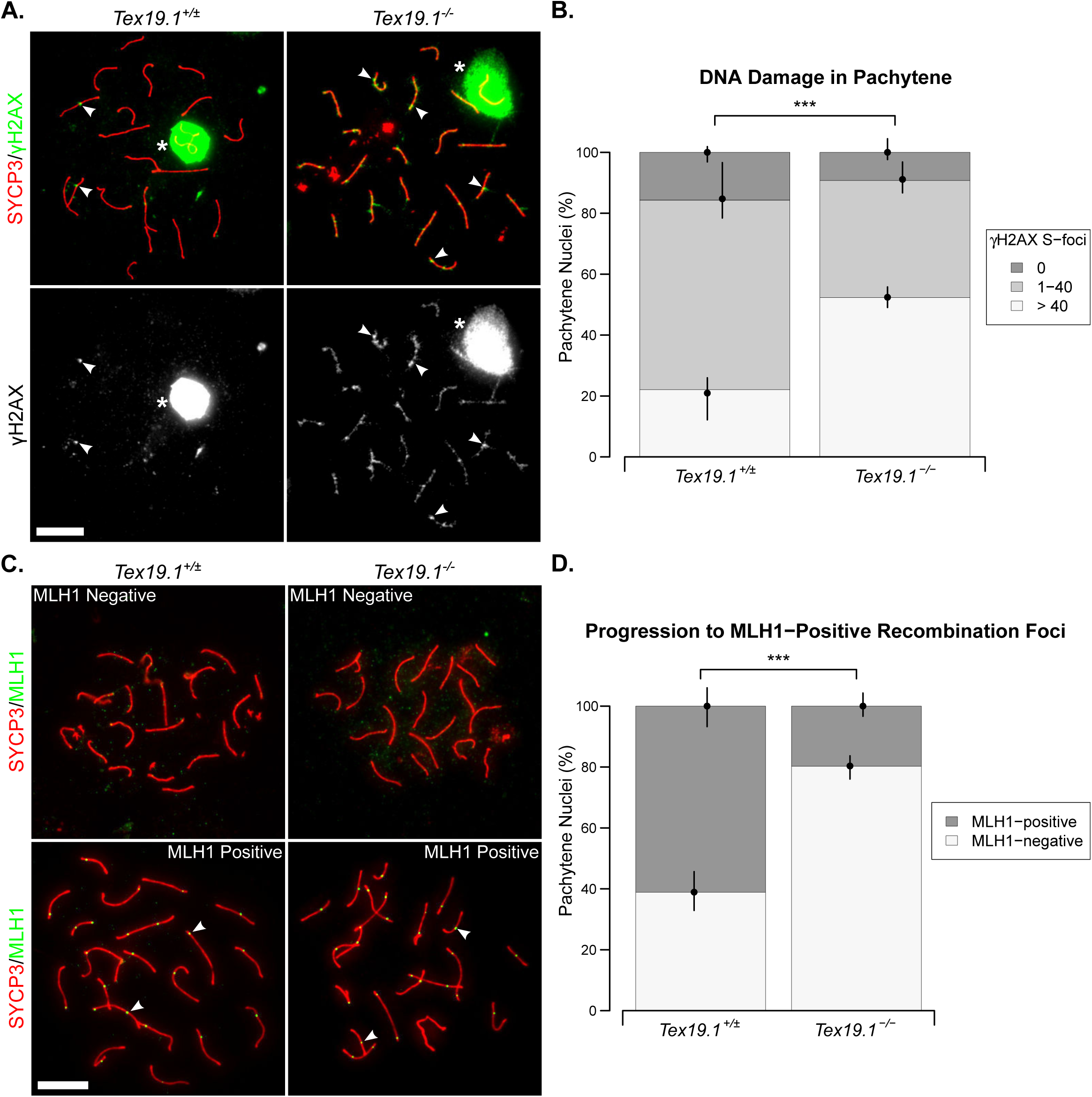
**Autosomally Synapsed Pachytene *Tex19.1***^***-/-***^**Spermatocytes Have Skewed Recombination Profiles.** (A) Immunostaining for γH2AX (green) in *Tex19.1*^*+/±*^ and *Tex19.1*^*-/-*^ autosomally synapsed pachytene spermatocyte chromosome spreads. SYCP3 (red) marks the lateral element of the synaptonemal complex. A single channel greyscale image for γH2AX is also shown. The *Tex19.1*^*+/±*^ image belongs to the intermediate γH2AX foci category (1-40 S-foci), the *Tex19.1*^*-/-*^ image belongs to the high γH2AX foci category (> 40 S-foci). Asterisks indicate sex bodies, arrowheads indicateexample S-foci associated with axes. Scale bars 10 μm. (B) Quantification of γH2AX S-foci. 172 and 120 autosomally synapsed *Tex19.1*^*+/±*^ and *Tex19.1*^*-/-*^ pachytene nuclei were categorised according to the number of γH2AX S-foci. 22.1%, 62.2%, and 15.7% of pachytene *Tex19.1*^*+/±*^ spermatocytes have > 40, 1-40 and 0 γH2AX foci respectively, compared to 52.3%, 38.5%, and 9.2% *Tex19.1*^*-/-*^ pachytene nuclei. Asterisks denote significance 
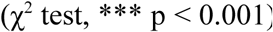
Nuclei were obtained from three animals for each genotype, and circles and vertical lines represent the means and interquartile distances of animals within each genotype. (C) Immunostaining for MLH1 (green) in control and *Tex19.1*^*-/-*^ autosomally synapsed pachytene spermatocyte chromosome spreads. SYCP3 (red) marks the lateral element of the synaptonemal complex. Example MLH1-positive and MLH1-negative images are shown. Nuclei containing ten or more axis-associated MLH1-foci (arrowheads) were classified as MLH1-positive. Scale bars 10 μm. (D) Quantification of pachytene nuclei that are MLH1-positive. 162 autosomally synapsed pachytene nuclei were scored for each genotype, 61.1% *Tex19.1*^*+/±*^ and 19.8% *Tex19.1*^*-/-*^ nuclei were MLH1-positive. Asterisks denote significance 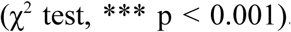Nuclei were obtained from three animals for each genotype, and circles and vertical lines represent the means and interquartile distances of animals within each genotype.

### Late Recombination Foci Are Depleted In Autosomally Synapsed *Tex19.1*^*-/-*^ Pachytene Spermatocytes

We next tested whether the increased frequency of early recombination intermediates and unrepaired DNA damage in pachytene *Tex19.1*^*-/-*^ spermatocytes is accompanied with any changes in the frequency of spermatocytes containing markers of late recombination foci. Autosomally synapsed pachytene *Tex19.1*^*-/-*^ spermatocytes were scored for the presence or absence of MLH1 (Figure 2C), a component of late recombination foci that marks crossover recombination events[29] 61% of control pachytene nuclei displayed MLH1 foci associated with autosomal axes, however this was reduced to just 20% among pachytene *Tex19.1*^*-/-*^ nuclei (Figure 2D). Thus, the population of autosomally synapsed pachytene *Tex19.1*^*-/-*^ spermatocytes has more early recombination foci, is enriched for nuclei containing large numbers of γH2AX S-foci, and is depleted for nuclei containing late recombination foci.

### Autosomally Synapsed *Tex19.1*^*-/-*^ Spermatocytes Establish Functional MSCI

The apparent enrichment of earlier substages in the population of pachytene *Tex19.1*^*-/-*^spermatocytes could reflect altered kinetics of recombination foci maturation during pachytene, or death of pachytene spermatocytes depleting later substages from the pachytene population. We have previously reported that loss of *Tex19.1* causes cell death in some spermatocytes [22], but this is an expected consequence of the synapsis defects, which can cause MSCI failure and apoptosis during pachytene [6,7]. However, the analyses performed in Figures 1 and 2 specifically selected pachytene nuclei that had no autosomal asynapsis. Cell death within the autosomally synapsed population ought to result in fewer cells progressing to diplotene, however the proportion of autosomally synapsed pachytene: diplotene spermatocytes is not significantly altered in *Tex19.1*^*-/-*^mice (Supporting Figure S2). Although the simplest interpretation of these data is that there is no significant cell death in autosomally synapsed *Tex19.1*^*-/-*^ pachytene spermatocytes, it is possible that MSCI is failing and causing cell death in a subset of fully synapsed pachytene *Tex19.1*^*-/-*^ cells, and that delays in pachytene progression in the surviving cells masks any change the autosomally synapsed pachytene: diplotene ratio. We therefore assessed whether MSCI is occurring normally in the absence of autosomal asynapsis in *Tex19.1*^*-/-*^ spermatocytes. HORMAD1 has a role in recruiting the transcriptional silencing machinery to asynapsed chromosomes [30], and defects in HORMAD dissociation from synapsed axes [11] could trigger MSCI failure in synapsed pachytene cells. However, HORMAD1 localisation is enriched on sex chromosome axes, and depleted from synapsed but not asynapsed autosomes as expected in *Tex19.1*^*-/-*^ spermatocytes (Figure 3A). Moreover, γH2AX immunostaining [27,31] suggests that the sex body itself is forming normally in autosomally synapsed *Tex19.1*^*-/-*^ synapsed pachytene nuclei (Figure 2A, asterisks). To test whether MSCI is being functionally established we immunostained meiotic chromosome spreads for RBMY, a Y-chromosome-encoded protein silenced by MSCI during pachytene [32]. RBMY was readily detected in asynapsed pachytene *Tex19.1*^*-/-*^ spermatocytes containing asynapsed autosomes but not in either control or *Tex19.1*^*-/-*^ autosomally synapsed pachytene spermatocytes (Figure 3B) suggesting that MSCI is occurring normally in autosomally synapsed spermatocytes in the absence of *Tex19.1*. Thus, autosomally synapsed *Tex19.1*^*-/-*^ pachytene spermatocytes do not have defects in MSCI that might trigger cell death in mid-late pachytene in the absence of asynapsis. Taken together, these data suggest that the enrichment of early recombination markers in autosomally synapsed *Tex19.1*^*-/-*^ spermatocytes likely represents progression through meiotic prophase being altered in at least some of these cells.

**Figure 3.**
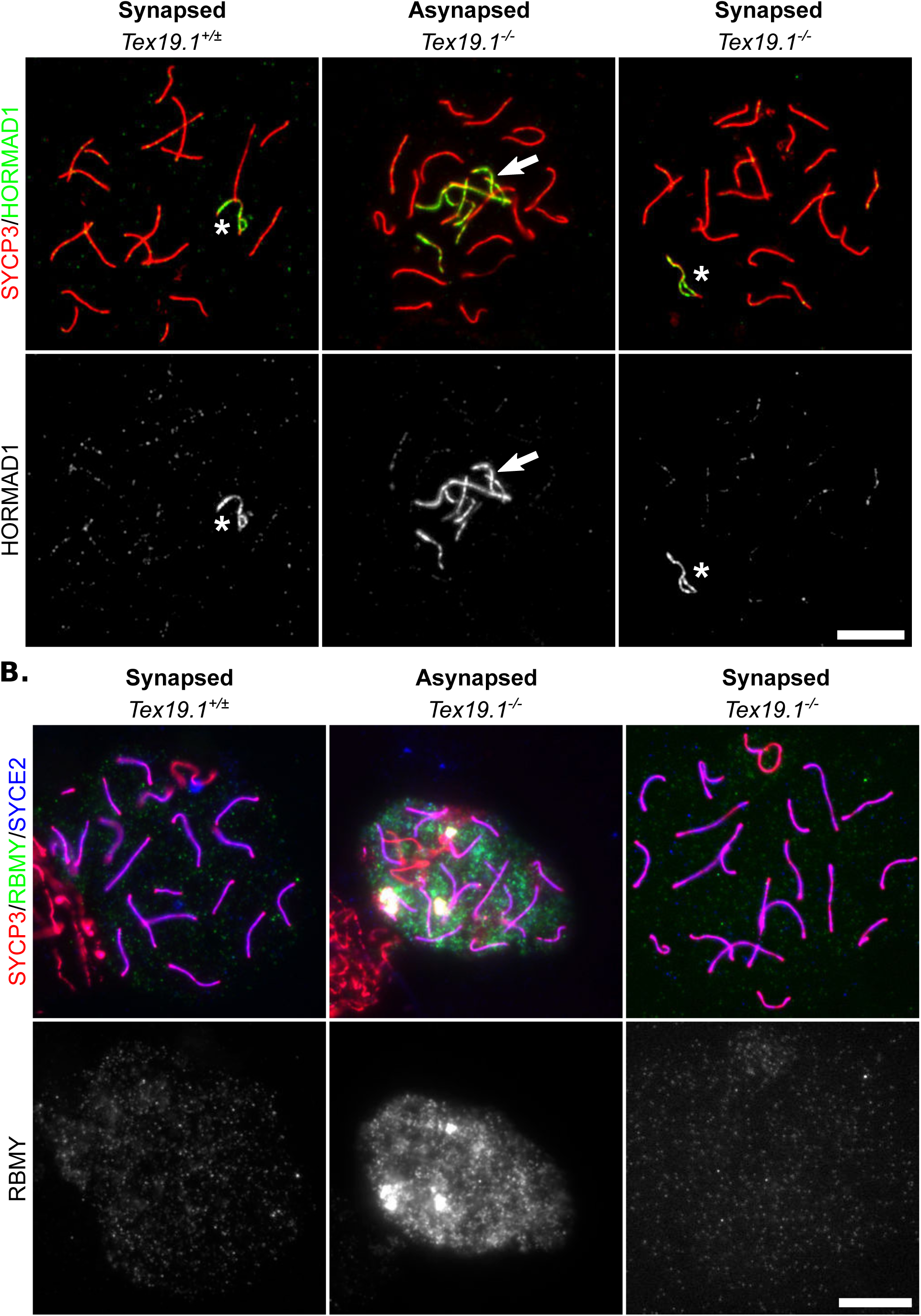
**Autosomally Synapsed Pachytene *Tex19.1^-/-^*Spermatocytes Correctly Localise HORMAD1 And Undergo Functional MSCI.** (A, B) Immunostaining for HORMAD1 (A, green) and RBMY (B, green) in *Tex19.1*^*+/±*^ and *Tex19.1*^*-/-*^ autosomally synapsed pachytene spermatocyte chromosome spreads. *Tex19.1*^*-/-*^autosomally asynapsed pachytene spermatocytes showing localisation of HORMAD1 to asynapsed autosomes, and failure to silence the Y-encoded RBMY protein are shown as positive controls. SYCP3 (A, B, red) and SYCE2 (A, blue) mark the lateral and central elements of the synaptonemal complex respectively. The sex body is marked with an asterisk. Single channel greyscale images for HORMAD1 and RBMY are also shown. Regions of adjacent non-pachytene nuclei are visible in some of the panels in B. RBMY images are representative of 64 *Tex19.1*^*+/±*^ nuclei and 78 *Tex19.1*^*-/-*^pachytene nuclei obtained from 3 animals of each genotype. HORMAD1 images are representative of 162 *Tex19.1*^*+/±*^ nuclei and 957 *Tex19.1*^*-/-*^ pachytene nuclei obtained from 3 *Tex19.1*^*+/±*^ and 3 *Tex19.1*^*-/-*^animals. Scale bars 10 μm.

### Pachytene Gene Expression And Histone Modifications Progress Abnormally In *Tex19.1*^*-/-*^ Spermatocytes

We next assessed whether the altered recombination kinetics in autosomally synapsed pachytene *Tex19.1*^*-/-*^ spermatocytes might be associated with more widespread changes in progression through pachytene. Expression of the testis-specific histone H1t is initiated in spermatocytes during mid-pachytene [25,33]. Histone H1t is expressed in *Dmc1*^*-/-*^ and *Msh5*^*-/-*^ asynapsed pachytene cells, suggesting that histone H1t expression does not depend on successful meiotic recombination or synapsis [6]. However, histone H1t is not expressed in arrested pachytene hypomorphic *Trip13*^*mod/mod*^ mutant spermatocytes, suggesting that it can be directly or indirectly affected by some perturbations in meiotic recombination [10]. Histone H1t staining is present throughout the chromatin of pachytene spermatocytes (Figure 4A), and while 68% of control pachytene spermatocytes express histone H1t, only 27% of autosomally synapsed pachytene *Tex19.1*^*-/-*^spermatocytes express this marker (Figure 4B). Thus, the abnormal recombination profiles seen in *Tex19.1*^*-/-*^ spermatocytes is accompanied by changes in the proportion of pachytene cells expressing a marker of mid-and late-pachytene spermatocytes.

**Figure 4.**
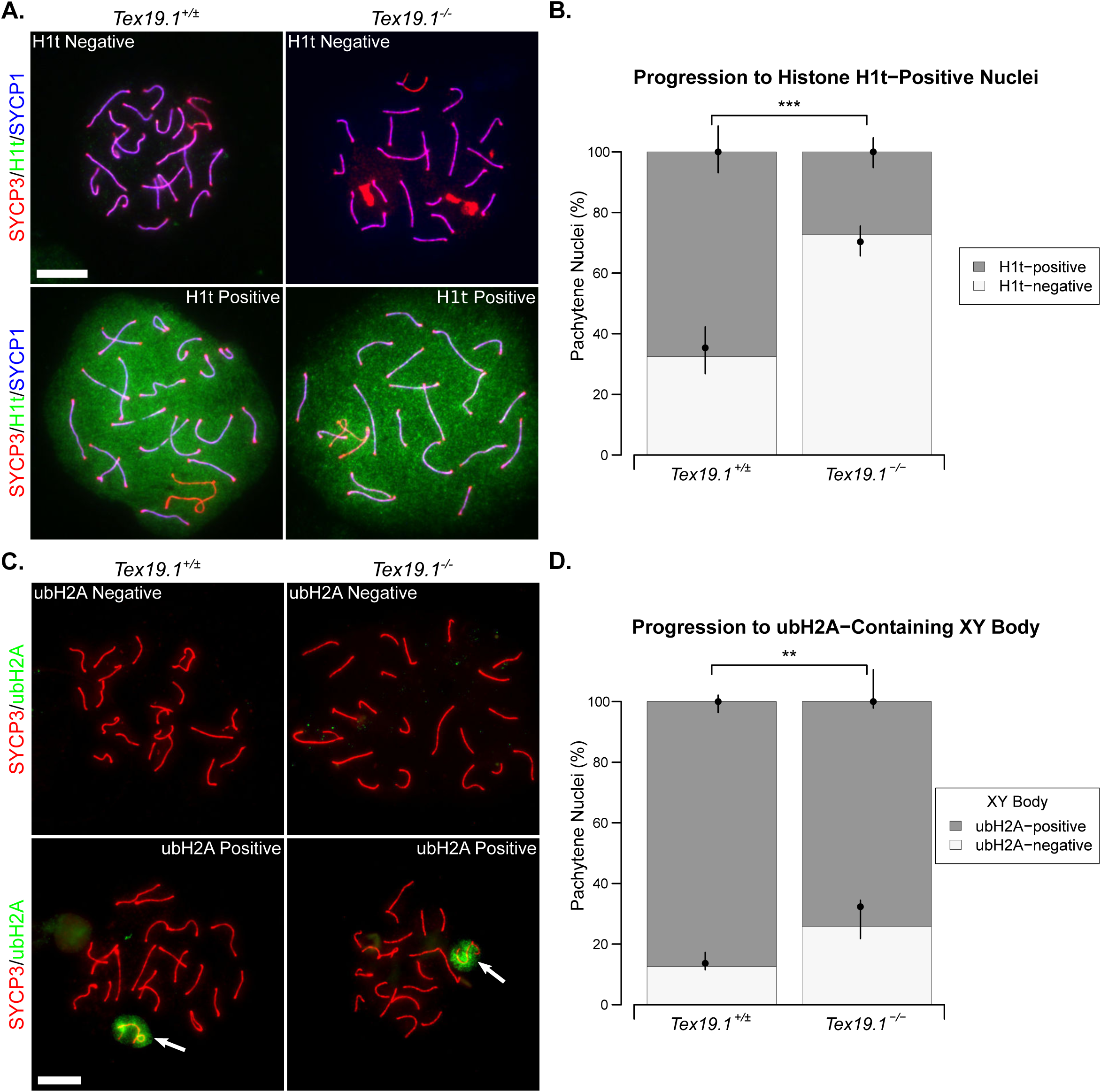
**Autosomally Synapsed Pachytene *Tex19.1^-/-^* Spermatocytes Have Delayed Progression Through Pachytene.** (A) Immunostaining for histone H1t (green) in *Tex19.1^+/±^* and *Tex19.1^-/-^* autosomally synapsed pachytene spermatocyte chromosome spreads. Example histone H1t positive and negative images are shown. SYCP3 (red) and SYCP1 (blue) mark the lateral elements and transverse filaments of the synaptonemal complex respectively. Scale bars 10 μm. (B) Quantification of the proportions of autosomally synapsed pachytene nuclei that are H1t-positive. 67.6% of *Tex19.1^+/±^* and 27.4% of *Tex19.1^-/-^* pachytene nuclei (n=111, 95; Fisher’s exact test, *** indicates p < 0.001) obtained from three animals of each genotype were H1t-positive. (C) Immunostaining for ubiquitylated H2A (ubH2A, green) in *Tex19.1^+/±^* and *Tex19.1^-/-^* autosomally synapsed pachytene spermatocyte chromosome spreads. Example ubH2A positive and negative images are shown. ubH2A clouds associating with the XY body are indicated by arrows. SYCP3 (red) marks the lateral element of the synaptonemal complex. Scale bars 10 μm. (D) Quantification of the proportions of autosomally synapsed pachytene nuclei that are have ubH2A clouds associated with the XY body. 87.3% of *Tex19.1^+/±^* and 74.1% of *Tex19.1^-/-^* pachytene nuclei (n=158, 147; Fisher’s exact test, ** indicates p < 0.01) obtained from four animals of each genotype have XY-associated ubH2A.

A number of changes in chromatin modifications occur as mouse spermatocytes progress through meiotic prophase [34], therefore we used these to assess pachytene progression in autosomally synapsed pachytene *Tex19.1*^*-/-*^ spermatocytes further. Mono-ubiquitylated histone H2A (ubH2A) accumulates at the sex body during pachytene, although it is not required for MSCI [35,36]. Immunostaining control spermatocytes for ubH2A revealed a cloud of staining at the sex chromosomes (Figure 4C), similar to that previously reported [35,36], in 87% of control pachytene spermatocytes (Figure 4D). A cloud of ubH2A staining was also detected at the sex chromosomes in autosomally synapsed *Tex19.1*^*-/-*^ pachytene spermatocytes (Figure 4C), however the proportion of cells with such staining was reduced to 74% (Figure 4D). Thus, the delay in maturation of recombination in autosomally synapsed *Tex19.1*^*-/-*^ pachytene spermatocytes is accompanied by more widespread changes in progression through pachytene.

In addition to the specific establishment of H2A mono-ubiquitylation at the sex body, the patterns of total ubiquitylation also vary as spermatocytes progress through pachytene [37,38]. Ubiquitylation can be broadly monitored using the FK2 monoclonal antibody which recognises both mono-and poly-ubiquitylation [39]. Therefore we investigated the progression of ubiquitylation in *Tex19.1*^*-/-*^ by immunostaining spermatocyte chromosome spreads with the FK2 antibody. Consistent with previous reports, FK2 staining in control pachytene spermatocytes is strikingly enriched at the sex chromosomes [37,38]. FK2 staining is initially restricted to the chromosome axes in the XY body in early pachytene, then extends throughout the XY body chromatin in mid and late pachytene [37,38]. Although both control and *Tex19.1*^*-/-*^ autosomally synapsed pachytene spermatocytes exhibit enriched FK2 staining on the sex chromosomes (Figure 5A), the frequency of pachytene nuclei with axial XY staining characteristic of early pachytene is higher in autosomally synapsed *Tex19.1*^*-/-*^spermatocytes (Figure 5B). Thus, similar to the recombination markers, γH2AX, histone H1t, and ubH2A analyses, sex chromosome ubiquitylation patterns suggest that autosomally synapsed *Tex19.1*^*-/-*^ spermatocytes are enriched for nuclei that display hallmarks of earlier substages of pachytene.

We also analysed the FK2 staining patterns on autosomes in pachytene *Tex19.1*^*-/-*^ spermatocytes. FK2 typically stains axial foci during early-mid pachytene, then becomes more diffusely localised throughout autosomal chromatin as pachytene progresses [38]. This accumulation of ubiquitylation on autosomal foci during early-mid pachytene potentially reflects the role of the ubiquitin proteasome system in maturation of recombination foci [38]. The frequency of autosomally synapsed pachytene nuclei with axial FK2 staining on autosomes (Figure 5C) is greatly increased in *Tex19.1*^*-/-*^ mice (Figure 5D). Therefore the altered patterns of ubiquitylation in *Tex19.1*^*-/-*^spermatocytes are not limited to ubH2A at the sex body, but extend to more general ubiquitylation at the sex chromosomes and autosomes. Similar to the altered recombination profiles and histone H1t marker analysis, the perturbed ubiquitylation patterns suggest that the population of autosomally synapsed *Tex19.1*^*-/-*^ spermatocytes is enriched for earlier substages of pachytene.

**Figure 5.**
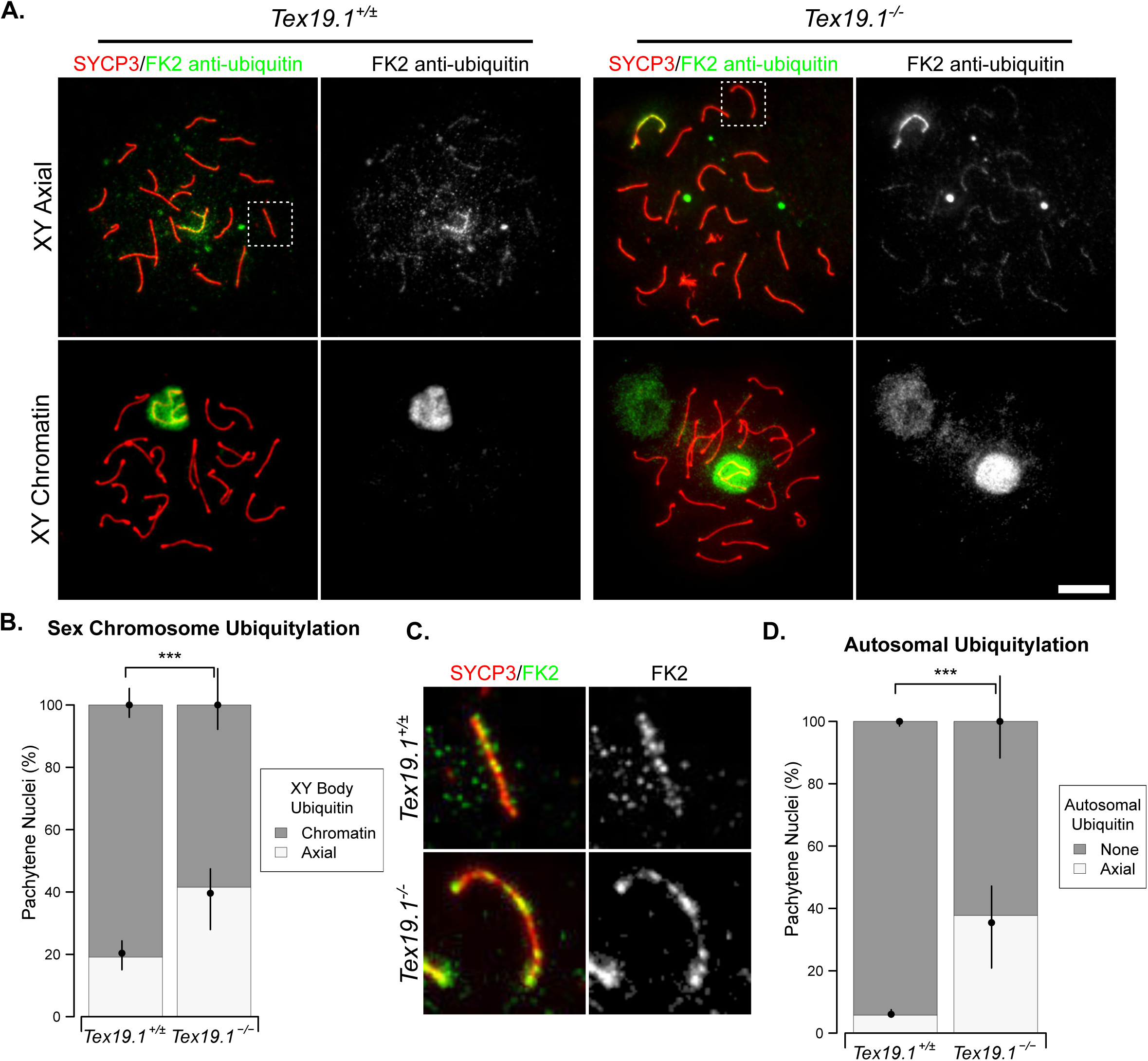
**Autosomally Synapsed Pachytene *Tex19.1*^*-/-*^ Spermatocytes Are Enriched For Ubiquitylation Patterns Associated With Early Pachytene.** (A) Immunostaining for mono-and poly-ubiquitylated proteins (FK2 anti-ubiquitin, green) in *Tex19.1*^*+/±*^and *Tex19.1*^*-/-*^autosomally synapsed pachytene spermatocyte chromosome spreads.Examples of nuclei with axial and chromatin staining on the XY body are shown. SYCP3 (red) marks the lateral element of the synaptonemal complex. A single channel greyscale image for FK2 is also shown. An adjacent non-meiotic nucleus positive for FK2 is visible in the XY chromatin *Tex19.1*^*-/-*^panel. Scale bars 10 μm. (B) Quantification of the proportions of autosomally synapsed pachytene nuclei with different sex chromosome ubiquitylation patterns. 19.1% of *Tex19.1*^*+/±*^ and 41.5% of *Tex19.1*^*-/-*^ pachytene nuclei obtained from three animals of each genotype have axial staining on the XY body 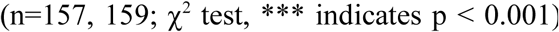(C) Higher magnification images of the boxed regions in panel A showing examples of FK2 staining on autosomal axes. A single channel greyscale image for FK2 is also shown. (D) Quantification of the proportions of autosomally synapsed pachytene nuclei with axial FK2 staining on their autosomes. 5.7% of *Tex19.1*^*+/±*^and 37.7% of *Tex19.1*^*-/-*^pachytene nuclei obtained from three animals of each genotype have FK2 staining on their autosomal axes 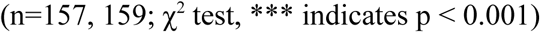

### *Tex19.1*^*-/-*^ Spermatocytes Have Reduced Chromosome Axis Elongation During Pachytene

The data on recombination and DNA damage markers, histone H1t and ubiquitylation together indicate that fully synapsed pachytene *Tex19.1*^*-/-*^ nuclei may be delayed in their progression from early to mid-late pachytene. However, it is not clear whether the autosomally synapsed *Tex19.1*^*-/-*^spermatocytes that reached late pachytene were still delayed in their meiotic progression. Progression through pachytene is reported to be associated with elongation of the autosomal chromosome axes between early and late pachytene [40], therefore we used chromosome axis length to assess whether late pachytene *Tex19.1*^*-/-*^ nuclei are still delayed in aspects of their meiotic progression. The late recombination marker MLH1 was used to select for spermatocytes in late pachytene for this analysis. Comparison of total axis length between *Tex19.1*^*+/±*^ and *Tex19.1*^*-/-*^MLH1-positive autosomally synapsed spermatocytes revealed a 6.5% reduction in length in the absence of *Tex19.1* (Figure 6A, Table 1). Comparison between individual and groups of axes ranked by size, as has been performed in similar analyses [13], demonstrated that all chromosome sizes were similarly affected (Figure 6B, Table 1). A significant reduction in length was not achieved for the smallest group of axes, though this is likely due to a greater degree of measurement error relative to total length. Thus, either elongation of the autosomal chromosome axes is delayed in fully synapsed late pachytene *Tex19.1*^*-/-*^ spermatocytes as part of a general perturbation of progression through pachytene in these mutants, or *Tex19.1*^*-/-*^ spermatocytes have shorter axes independent of the other changes in progression through pachytene.

**Figure 6.**
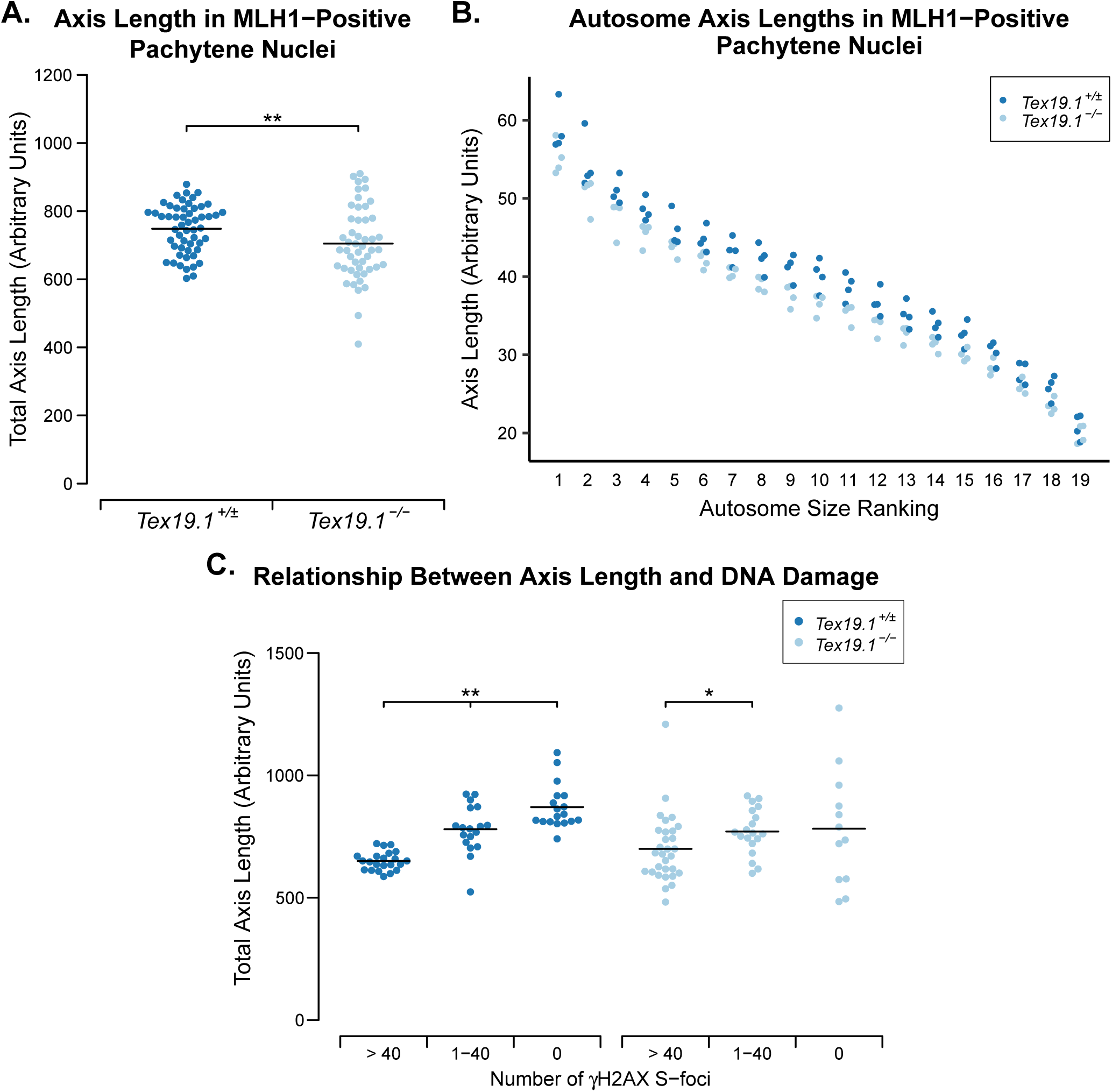
**Axis Elongation During Pachytene Is Perturbed In Autosomally Synapsed Pachytene *Tex19.1***^***-/-***^**Spermatocytes.** (A) Quantification of chromosome axis length in MLH1-positive autosomally synapsed *Tex19.1*^*+/±*^ and *Tex19.1*^*-/-*^ pachytene spermatocytes. Axis length was measured from SYCP3 immunostaining of chromosome spreads. The total length of all nineteen autosomes for individual nuclei is shown, and the mean values for each genotype (748±10 and 705±15 arbitrary units) indicated with a horizontal line. A total of 54 and 49 nuclei from four *Tex19.1*^*+/±*^ and *Tex19.1*^*-/-*^ animals respectively werescored, ** indicates p < 0.01 (Mann-Whitney U test). (B) Axis lengths of individual size-ranked autosomes from MLH1-positive autosomally synapsed *Tex19.1*^*+/±*^ and *Tex19.1*^*-/-*^ pachytene spermatocytes shown in panel A. (C) Plot showing relationship between γH2AX S-foci and axis elongation in autosomally synapsed *Tex19.1*^*+/±*^ and *Tex19.1*^*-/-*^ pachytene spermatocytes. Total axis lengths for individual nuclei belonging to the indicated categories of γH2AX S-foci are plotted. In contrast to panel A, these nuclei were not selected for being MLH1-positive. Means for each group are indicated by horizontal lines (1032±13, 1237±37, 1382±34, 1110±40, 1223±34, 1285.7±74.3 from left to right, n=22, 18, 18, 30, 19, 18 respectively). Statistically significant differences between groups were determined by Mann-Whitney U test, ** indicates p < 0.01, * indicates p < 0.05. None of the comparisons between the *Tex19.1*^*+/±*^ and *Tex19.1*^*-/-*^ groups are significantly different. Nuclei were derived from three control animals and four *Tex19.1*^*-/-*^ animals.

**Table 1.** Axis Lengths Of Groups Of Size-Ranked Autosomes In MLH1-Positive Autosomally Synapsed Pachytene Nuclei.

| Grouped<br>Axis Ranks | Mean group axis length (arbitrary units) |  | Difference<br>in *Tex19.1*<sup>-/-</sup> |
| --- | --- | --- | --- |
|  | *Tex19.1*<sup>+/-</sup> | *Tex19.1*<sup>-/-</sup> |  |
| **1-2** | 113.69 | 105.79 | -6.9% * |
| **3-5** | 146.20 | 137.04 | -6.3% * |
| **6-11** | 248.31 | 231.90 | -6.6% * |
| **12-16** | 166.92 | 155.49 | -6.8% * |
| **17-19** | 73.53 | 69.80 | -5.1% |
| **Total** | 748.64 | 700.01 | -6.5% ** |

Autosomes were ranked by size and grouped as indicated. The mean total axis lengths for each group of autosomes is shown. Four animals were analysed for each genotype and a total of 50 and 55 nuclei analysed for *Tex19.1*^*+/±*^ and *Tex19.1*^*-/-*^ respectively. Student’s t-test was used to test for statistical signficance between genotypes (* indicates p < 0.05, ** indicates p < 0.01.)

To investigate whether loss of *Tex19.1* results in shorter chromosome axes independently of altered progression through pachytene we used the repair of γH2AX S-foci to monitor progression through pachytene. *Tex19.1*^*+/±*^ and *Tex19.1*^*-/-*^ spermatocytes both elongated their chromosome axes as they progressed through pachytene and repaired γH2AX S-foci (Figure 6C). However, when autosomally synapsed *Tex19.1*^*+/±*^ and *Tex19.1*^*-/-*^ pachytene spermatocytes with similar amounts of DNA damage are compared, no significant differences in axis length are evident (Figure 6C). Therefore, the shorter chromosome axes in MLH1-positive *Tex19.1*^*-/-*^ pachytene spermatocytes likely reflects the perturbed progression of these spermatocytes through pachytene, and axis length appears to be more closely associated with repair of γH2AX S-foci than appearance of MLH1 foci during pachytene.

### Delayed Progression Through Pachytene in *Tex19.1*^*-/-*^ Spermatocytes is Dependent on *Spo11*

Taken together, the data presented in this manuscript suggests that multiple aspects of progression to late pachytene are delayed in autosomally synapsed *Tex19.1*^*-/-*^ mutants. This phenotype could potentially reflect the delayed accumulation of early recombination generated in zygotene *Tex19.1*^*-/-*^spermatocytes and the activity of a checkpoint co-ordinating progression through pachytene with repair of DNA damage or maturation of recombination foci. Alternatively, the altered progression through pachytene in autosomally synapsed *Tex19.1*^*-/-*^ spermatocytes could potentially reflect a recombination-independent role for *Tex19.1* in co-ordinating pachytene. To distinguish these possibilities we assessed progression through pachytene in *Tex19.1*^*-/-*^ spermatocytes in the absence of meiotic DSBs. *Spo11*^*-/-*^ spermatocytes fail to generate meiotic DSBs and as such are unable to undergo meiotic recombination and arrest in a zygotene-like state with defective chromosome synapsis [41,42]. Despite this, *Spo11*^*-/-*^ spermatocytes are able to progress to a mid-pachytene-like state of gene expression and recruit histone H1t [43]. Thus, we could test whether the delay in histone H1t expression seen in pachytene *Tex19.1-/-* spermatocytes depends on the presence of *Spo11* and meiotic recombination. Consistent with a previous report [43] we detected H1t staining in approximately 50% of zygotene-like control *Spo11*^*-/-*^ mutant spermatocytes (Figure 7A, Figure 7B). In *Tex19.1*^*-/-*^*Spo11*^*-/-*^ double knockout mice a similar frequency of zygotene-like spermatocytes were H1t-positive (Figure 7A, Figure 7B). Indeed, the proportion of histone H1t-positive zygotene-like cells in *Tex19.1*^*-/-*^*Spo11*^*-/-*^ double mutant mice is higher than the proportion of H1t-positive autosomally synapsed cells in *Tex19.1*^*-/-*^ single mutant mice (Figure 4B, Figure 7B). Therefore the reduced progression of *Tex19.1*^*-/-*^ spermatocytes to an H1t-positive mid-pachytene-like state of gene expression is dependent on the meiotic DSB forming endonuclease *Spo11*. This finding indicates that the delayed progression through pachytene observed in *Tex19.1*^*-/-*^ spermatocytes is related to the meiotic recombination defects incurred in this mutant and is consistent with the presence of a recombination-dependent checkpoint regulating progression through pachytene.

**Figure 7.**
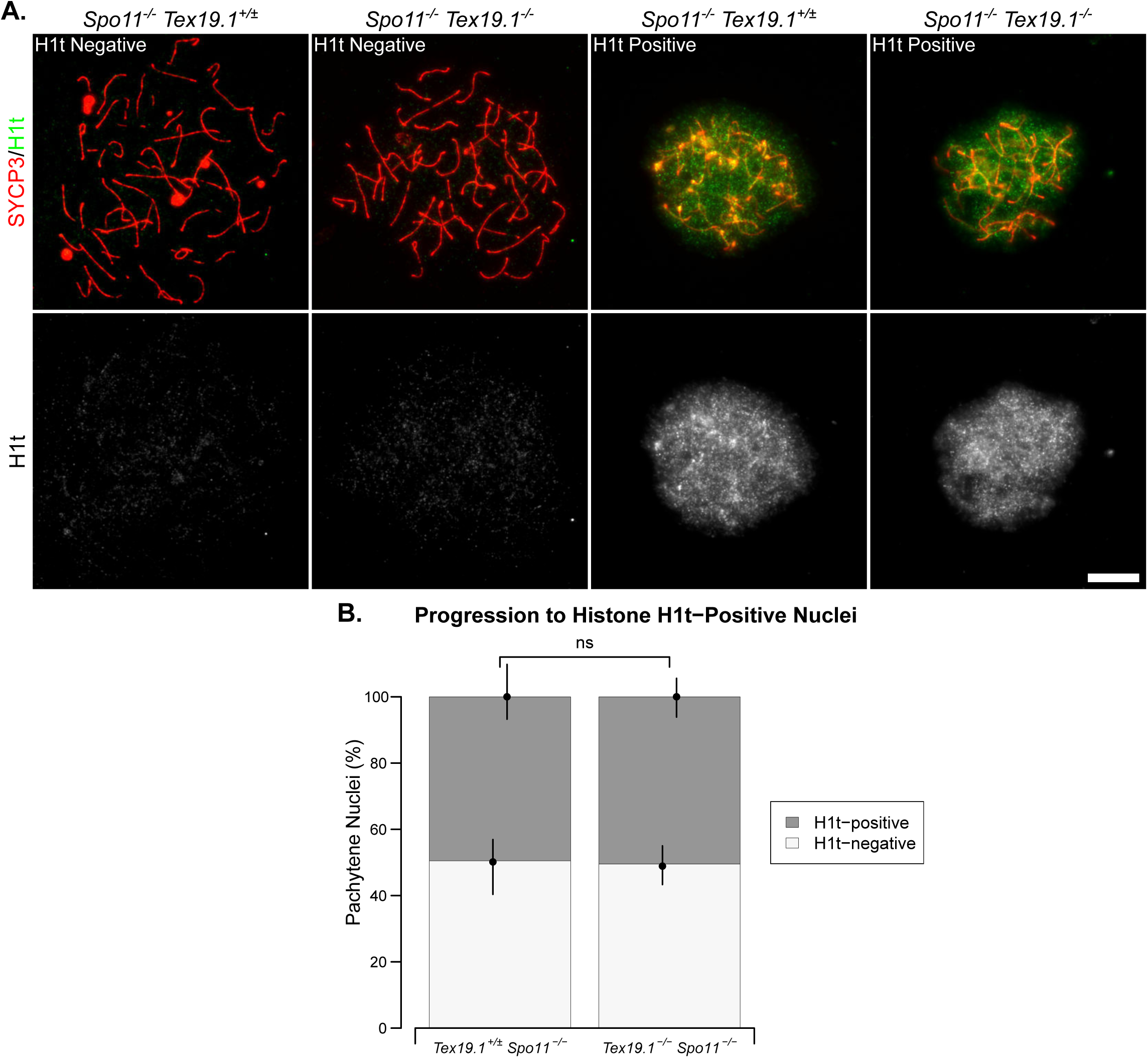
**Delayed Histone H1t Expression In *Tex19.1***^***-/-***^**Spermatocytes Depends On *Spo11*.** (A) Immunostaining for histone H1t (green) in *Spo11*^*-/-*^*Tex19.1*^*+/±*^ single knock-out and *Spo11*^*-/-*^*Tex19.1*^*-/-*^double knockout zygotene-like spermatocyte chromosome spreads. Example histone H1tpositive and negative images are shown. SYCP3 (red) marks the lateral element of the synaptonemal complex respectively. Scale bars 10 μm. (B) Quantification of the proportions of zygotene-like nuclei that are H1t-positive. 49.5% of *Spo11*^*-/-*^*Tex19.1*^*+/±*^ and 50.5% of *Spo11*^*-/-*^*Tex19.1*^*-/-*^zygotene-like nuclei obtained from three animals of each genotype were H1t-positive (n=97, 99; χ^2^ test, ns indicates no significant difference, p > 0.05).

## Discussion

In this study we have investigated whether delayed meiotic recombination has consequences for progression through pachytene in mouse spermatocytes. The data presented in this study show that autosomally synapsed *Tex19.1*^*-/-*^ spermatocytes have perturbed progression through pachytene (Figure 8) and that at least some aspects of this altered progression depend on *Spo11*. These data suggest that *Tex19.1*^*-/-*^ spermatocytes are activating a recombination-dependent checkpoint that coordinately regulates aspects of progression through pachytene. The skewed maturation of recombination markers in pachytene *Tex19.1*^*-/-*^ spermatocytes potentially reflects substrate-product relationships between the delayed accumulation of early recombination foci in zygotene and their subsequent maturation. Indeed, mouse spermatocytes with altered *Spo11* expression have a delayed burst of DSB formation in zygotene that also translates into an enrichment for early recombination markers during pachytene [44]. In addition, the close association between axis length and repair of DNA damage during pachytene could potentially also reflect a causal relationship between these parameters. However, other aspects of the skewed progression of *Tex19.1*^*-/-*^ spermatocytes, such as the delay in expression of histone H1t, would seem more likely to represent activation of a checkpoint co-ordinately regulating independent meiotic processes. This recombination-dependent checkpoint appears to be acting in early pachytene and delaying, but not preventing, progression to late pachytene (Figure 8).

**Figure 8.**
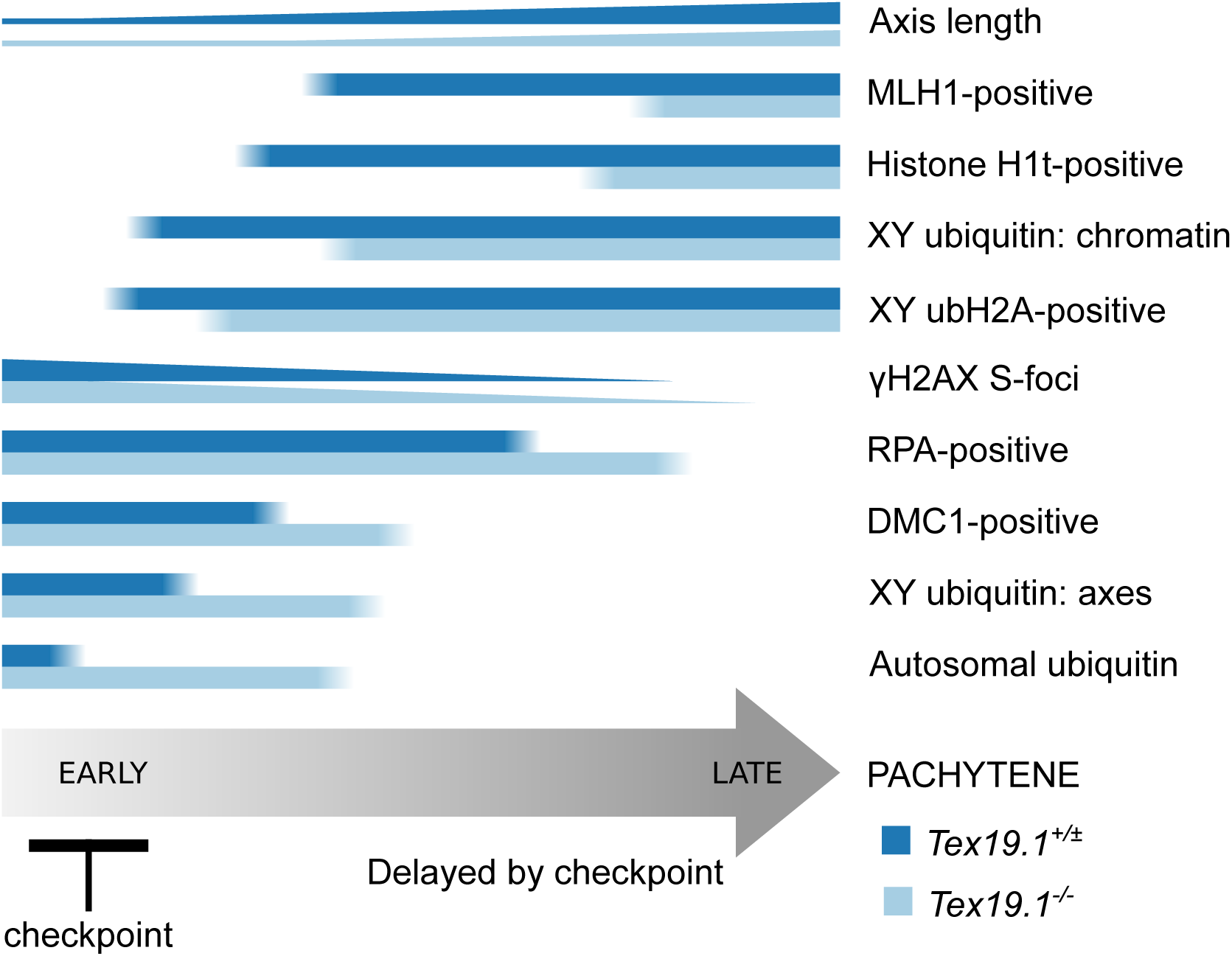
**Schematic Diagram Showing Delayed Progression Through Pachytene In Autosomally Synapsed Pachytene *Tex19.1*^*-/-*^ Spermatocytes**. Summary of aspects of progression through pachytene analysed in this study. Horizontal bars indicate the proportion of pachytene nuclei expressing different markers of progression throughpachytene. Number of γH2AX foci and length of axis are indicated by the height of the bars. Proportions of DMC1 and RPA positive nuclei are estimated from the violin plots in Supporting Figure S1. The differences in the behaviour of these markers in autosomally synapsed pachytene *Tex19.1*^*+/±*^and *Tex19.1*^*-/-*^spermatocytes is indicated by the blue bars. *Tex19.1*^*-/-*^spermatocytes appear to activate a *Spo11*-dependent recombination checkpoint in early pachytene that delays progression to late pachytene.

Although we have shown that delayed progression through pachytene in *Tex19.1*^*-/-*^ spermatocytes depends on *Spo11*, it is not clear what recombination-dependent lesion is activating this checkpoint. Normal recombination intermediates could have deleterious effects if they persist during the meiotic nuclear divisions [45], and early recombination intermediates could be activating a checkpoint in pachytene *Tex19.1*^*-/-*^ spermatocytes. This possibility implies that the checkpoint may have a role in co-ordinating progression through pachytene during normal meiosis in wild-type cells by delaying aspects of progression through pachytene until recombination proceeds beyond a certain point. Alternatively, *Tex19.1*^*-/-*^ spermatocytes could have additional defects in processing recombination intermediates and abnormal lesions arising during maturation of recombination intermediates could be activating this checkpoint. Moderate hypomorphic mutations in *Trip13* cause defects in dissociation of HORMAD1 from synapsed autosome axes, defects in processing recombination intermediates, and activation of a checkpoint that prevents expression of histone H1t in spermatocytes [10,,–13]. *Tex19.1*^*-/-*^ spermatocytes do not phenocopy *Trip13* mutants with respect to HORMAD1 dissociation, and if loss of *Tex19.1* is causing similar defects in maturation of recombination then, in contrast to *Trip13* mutants, these lesions are not severe or persistent enough to trigger cell death. Spermatocytes with altered expression of *Spo11* exhibit a delayed burst of DSB formation and completely synapse their autosomes [44], and analysis of histone H1t expression in these spermatocytes may help to distinguish whether delays in early meiotic recombination are sufficient to trigger the pachytene checkpoint described here. In addition, the *Trip13* checkpoint arrest is mediated by CHK2, a component of the ATM-dependent DNA damage signalling cascade. Investigating the role of CHK2 and ATM signalling in the *Tex19.1*^*-/-*^ checkpoint delay may help determine whether these mutants are activating the same checkpoint with different outcomes.

The data presented in this study provides evidence that a recombination checkpoint operating in mouse spermatocytes can act to delay progression through pachytene. Although there are examples of checkpoints inducing pachytene delays in yeast [46], worms [47] and flies [48], the synapsis and recombination checkpoints operating in pachytene mouse spermatocytes have primarily been shown to trigger cell death when activated and act as a quality control [7,10]. This study extends these findings and suggests that checkpoint responses can regulate and co-ordinate progression of mouse spermatocytes through pachytene. The change in progression through pachytene in response to delayed recombination that we have identified would potentially allow or reflect the repair of recombination-dependent lesions in these cells before they continue further in the meiosis.

## Materials and Methods

### Mice

*Tex19.1*^*-/-*^ animals on a C57BL/6 genetic background were bred and genotyped as described [22]. *Spo11*^*+/-*^ heterozygous mice [41] on a C57BL/6 genetic background [6] were inter-crossed with *Tex19.1*^*+/-*^ mice. Animal experiments were carried out under UK Home Office Project Licence PPL 60/4424. Animals were culled by cervical dislocation. *Tex19.1*^*+/+*^ and *Tex19.1*^*+/-*^ animals have no difference in spermatogenesis as assessed by epididymal sperm counts [22] and data from these control genotypes were pooled as *Tex19.1*^*+/±*^.

### Immunostaining Meiotic Chromosome Spreads

Chromosome spreads were prepared as described by Peters et al. [49], or by Costa et al. [50] and stored at -80ºC until use. After thawing, chromosome spreads were blocked with PBS containing 0.15% BSA, 0.1% Tween-20 and 5% goat serum. Primary antibodies were also diluted in PBS containing 0.15% BSA, 0.1% Tween-20 and 5% goat serum as indicated in Supplementary Table 1. Alexa Fluor-conjugated secondary antibodies (Invitrogen) were used at a 1:500 dilution, and DNA was stained by including 2 ng/μl 4,6-diamidino-2-phenylidole (DAPI) in the secondary antibody incubations. Slides were mounted in 90% glycerol, 10% PBS, 0.1% p-phenylenediamine, and images captured with iVision or IPLab software (BioVision Technologies) using an Axioplan II fluorescence microscope (Carl Zeiss) equipped with motorised colour filters. At least three experimental and three control animals were analysed for each experiment. RAD51 and RPA recombination foci were imaged by capturing z-stacks using a piezoelectrically-driven objective mount (Physik Instrumente) controlled with Volocity software (PerkinElmer). These images were deconvolved using Volocity, and a 2D image generated in Fiji [51].

Nuclei were staged by immunostaining for the axial/lateral element marker SYCP3 [52]. Pachytene nuclei were identified in this analysis by complete co-localisation of SYCP3 and either the transverse filament marker SYCP1 [53] or the central element marker SYCE2 [50] on all nineteen autosomes, or by the presence of nineteen bold SYCP3 axes in addition to two paired or unpaired sex chromosome axes of unequal length. The synapsis status of the X and Y chromosomes was not considered in these analyses as the timing and kinetics of sex chromosome synapsis differs from autosomes [54]. Asynapsed pachytene *Tex19.1*^*-/-*^ nuclei [22] were distinguished from zygotene due to incomplete sets of synapsed autosomes. Zygotene-like nuclei in *Spo11*^*-/-*^ mice were identified by the presence of fully formed axial elements.

DMC1, RAD51, RPA and MLH1 foci were counted as recombination foci when they overlapped a chromosome axis, and the foci counts reported refer to autosomal foci only. Nuclei were classed as MLH1-positive if they contained more than ten axial MLH1 foci. For γH2AX foci, prominent foci associated with SYCP3-stained axes were scored. For RBMY and histone H1t scoring, nuclei were imaged with fixed exposures, the image filenames were then randomised by computer script and scored blind as positive or negative. For analysis of FK2 staining, axial autosome staining is significantly less intense than the signal in the XY body, and image contrast was adjusted appropriately to score each of these parameters. Axis length was measured using NeuronJ [55] as described previously [56]. All scoring of immunostained chromosome spreads was performed blind on images after randomisation of filenames by computer script. Data were analysed in R [57], means are reported ± standard error. Nuclei were typically obtained from at least three animals for each genotype; for categorical data, stacked bar graphs represent proportions of all nuclei scored, while the means and interquartile ranges indicate the variation between animals of the same genotype.

## Acknowledgements

We also thank C. James Ingles (University of Toronto, Canada) for anti-RPA antibodies, David Elliott (University of Newcastle, UK) for anti-RBMY antibodies, Mary Ann Handel (Jackson Laboratory, Bar Harbor, USA) for anti-histone H1t antibodies, Attila Tóth (Technische Universität Dresden, Germany) for anti-HORMAD1 antibodies, Howard Cooke (MRC HGU, Edinburgh, UK) for anti-SYCE2 antibodies, and James Turner (Crick Institute, London, UK) and Bernard de Massy (IGH, Montpellier, France) for *Spo11* mutant mice. We thank the animal and imaging facility staff at MRC HGU for their help and advice. We thank Chris Playfoot and Issy MacGregor for comments on the manuscript, and other members of the Adams lab and MRC HGU for advice and discussion.

